# Transcranial photoacoustic characterization of neurovascular physiology during early-stage photothrombotic stroke in neonatal piglets *in vivo*

**DOI:** 10.1101/2021.07.08.451613

**Authors:** Jeeun Kang, Xiuyun Liu, Suyi Cao, Steven R. Zeiler, Ernest M. Graham, Emad M. Boctor, Raymond C. Koehler

**Author notes:** Corresponding authors: Emad M. Boctor, Ph.D. Raymond C. Koehler, Ph.D. These authors equally contributed (Jeeun Kang, Xiuyun Liu).

## Abstract

Perinatal ischemic stroke is estimated to occur in 1/2300–1/5000 live births, but early differential diagnosis from global hypoxia-ischemia is often difficult. In this study, we tested the ability of a hand-held transcranial photoacoustic (PA) imaging to non-invasively detect a focal photothrombotic stroke (PTS) within 2 hours of stroke onset in a gyrencephalic piglet brain. 17 stroke lesions of approximately 1-cm^2^ area were introduced randomly in anterior or posterior cortex via the light/dye PTS technique in anesthetized neonatal piglets (*n* = 11). The contralateral non-ischemic region served as control tissue for discrimination contrast for the PA hemoglobin metrics: HbO_2_ saturation, total hemoglobin (tHb), and individual quantities of oxygenated and deoxygenated hemoglobin (HbO_2_ and HbR). The PA-derived tissue HbO_2_ saturation at 2 hours yielded a significant separation between control and affected regions-of-interest (*p* < 0.0001), which were well matched with 24-hr post-stroke cerebral infarction confirmed in the triphenyltetrazolium chloride (TTC)-stained image. The quantity of HbO_2_ also displayed a significant contrast (*p* = 0.021), whereas tHb and HbR did not. The analysis on receiver operating characteristic curves and multivariate data analysis also agreed with the results above. This study shows that a hand-held transcranial PA neuroimaging can detect a regional thrombotic stroke in cerebral cortex of a neonatal piglet. In particular, we conclude that the HbO2 saturation metric can be used alone to identify regional stroke lesions. The lack of change in tHb may be related to arbitrary hand-held imaging configuration and/or entrapment of red blood cells within the thrombotic stroke.

## Introduction

Perinatal arterial ischemic stroke is estimated to occur in 1/2300–1/5000 live births, and can result in long-term deficits in motor, cognitive, attention, and executive functions and persistent seizures [1–4]. Despite of all the efforts, there is still no effective treatment available to improve the functional recovery after perinatal stroke, and “damage control” based on a rapid detection and following neuroprotective treatment has been the best strategy in clinical use [5,6]. The traditional strategy is mainly based on a supportive care [7,8]. Several fetal monitoring technologies have been developed to enable the prompt treatment but have great limitations. Electronic fetal heart rate monitoring (EFHM) has been a standard of care in perinatal monitoring since 1970s. However, no benefit has been shown in l cerebral palsy by using EFHM, while the cesarean delivery has increased from 5% to more than 30% since using this technology [9]. Meanwhile, several studies showed that EFHM has a critical false-positive rate, even as high as 99.8 % [10], and not sensitive for perinatal stroke detection in fetal brain [11]. Magnetic resonance imaging (MRI) is the definitive diagnostic tool for ischemic stroke after birth, but its application in the newborn is questionable due to the inaccessibility in the first day after birth [12]. As alternatives, several biophotonic modalities have been proposed to allow cost-effective and continuous monitoring of neonatal brain in this period. Near-infrared spectroscopy (NIRS) and diffused optical tomography (DOT) have shown their capability of non-invasive, continuous, and fast monitoring of cerebral oxygenation level and hemodynamic changes in the neonatal brain [13,14]. However, both methods suffer from low spatial specificity and incapability of distinguishing the arterial and venous compartments, leading to low clinical specificity. Therefore, there is a critical need for effective and prompt monitoring of the perinatal brain preferably with quantitative metrics of physiological dynamics to enable early-stage decision and immediate commencement of the treatments.

Photoacoustic (PA) sensing modality, a combination of ultrasound and optical modalities, has been highlighted recently due to its high spatiotemporal resolution and deep imaging depth, which may play an alternative role for the unmet clinical need. The technology emits the pulse laser light on intact skin, and the light penetrates into deep tissue determined by the distribution of tissue scattering coefficients. After being absorbed at localized tissue with unique absorption coefficients, the conversion of absorbed energy to instantaneous heat would generate the thermal elastic expansion, which leads to pressure which is detected by an external ultrasound transducer [15]. Several clinical applications using PA technology have been proposed [16–20], and hemodynamic imaging has been a primary clinical application with stark hemoglobin absorbance, enabling quantifications of blood oxygen saturation and cerebral blood volume [21–23]. Transcranial PA sensing of physiological change at the superior sagittal sinus (SSS) has been demonstrated in the neonatal piglet and ovine models through intact scalp [24–26], but there is still room for further development to have physiological measures in brain tissue, wherein the concentration of hemoglobin is an order of magnitude less than in the sagittal sinus vein. J. Lv, *et al.* also demonstrated transcranial PA imaging of structural and functional dynamics over rodent brain during ischemic stroke at a very early stage (from 5-min to 6-hour onset) [27], but the implication for clinical translation is still limited by the use of the rodents with impractical imaging scale and small signal attenuation with its very thin skull layer.

Therefore, development of a preclinical stroke model that has a scale and maturity level comparable to human newborns is an important component for the evaluation of a novel fetal brain monitoring modality. We chose to study newborn piglets with a 40 g gyrencephalic brain in which neuroanatomical and neurophysiological features well resembling those in term human infants can be achieved [28–30] with similar brain growth and developmental patterns [30–32]. A variety of adult rodent ischemic stroke models have been developed, including internal carotid arterial suture occlusion of the middle cerebral artery (MCAO), direct surgical occlusion through a craniotomy, direct application of endothelin-1 on the vessels, injection of an embolus, and photothrombosis in the light/dye model. However, because the piglet has a rete mirablis, the internal carotid arterial suture model and embolic model are not feasible [33]. On the other hand, the photothrombosis (PTS) stroke model induces ischemia in a well-defined region, with minimal surgical intervention, low mortality, and high reproducibility [34]. The PTS technique induces a cortical infarction through the photo-activation of a light-sensitive dye (e.g., Rose bengal, erythrosin B) previously delivered into the circulation, resulting in local vessel thrombosis in the areas exposed to the light [35]. This model has been used successfully in the newborn piglet [28,29]. When the circulating dye is illuminated at the effective wavelength, it generates free radicals that lead to endothelial damage, platelet activation, and thrombosis in both pial and intraparenchymal vessels within the irradiated area [34].

Herein, we present transcranial PA imaging of regional ischemic stroke in the neonatal piglet model. We specifically assumed PA imaging metrics that can be achieved in a bedside diagnostic setup using a hand-held PA imaging device in order to achieve a critical translational milestone towards its clinical application. There are three aims of this study: (1) To establish stable regional stroke models in the neonatal piglet; (2) to investigate whether ischemic features in the PA imaging, such as oxyhemoglobin (HbO_2_) saturation, total hemoglobin concentration (tHb), and HbO_2_ and HbR quantities, can contribute to an accurate localization of PTS lesions; (3) to validate whether those early features obtained in 2 hours post-stroke period in a hand-held PA imaging form factor can predict cerebral infarction at a later time point.

## Results

### Photothrombotic stroke *in vivo*

Figure 1 shows the experimental setup to induce 17 focal infarctions in 11 piglets (weight 1.35 kg – 1.80 kg). In detail, 5 piglets received one PTS lesion for the controlled characteristics of PA imaging PTS lesions, while the other 6 piglets received two lesions to mimic the unpredictable nature of stroke induction more accurately. For the latter group, the lesions were randomly selected among pre-determined four candidate positions at frontal and posterior cortex regions (Figure 1c). The single-lesion group only considered the posterior cortex regions. We anticipated no difference in the generation of PTS lesions depending on the cortical position, following the mechanism of PTS protocol generating thrombosis in direct region of light illumination. The cortical infarction width (lateral) and length (sagittal) was identified by contouring the white surface area in TTC-stained brain. The PTS protocol yielded well controlled mean cortical infarction area of 1 cm^2^ with 0.89 ± 0.20 cm and 1.03 ± 0.41 cm of width and length (mean ± SD), respectively.

**Figure 1.**
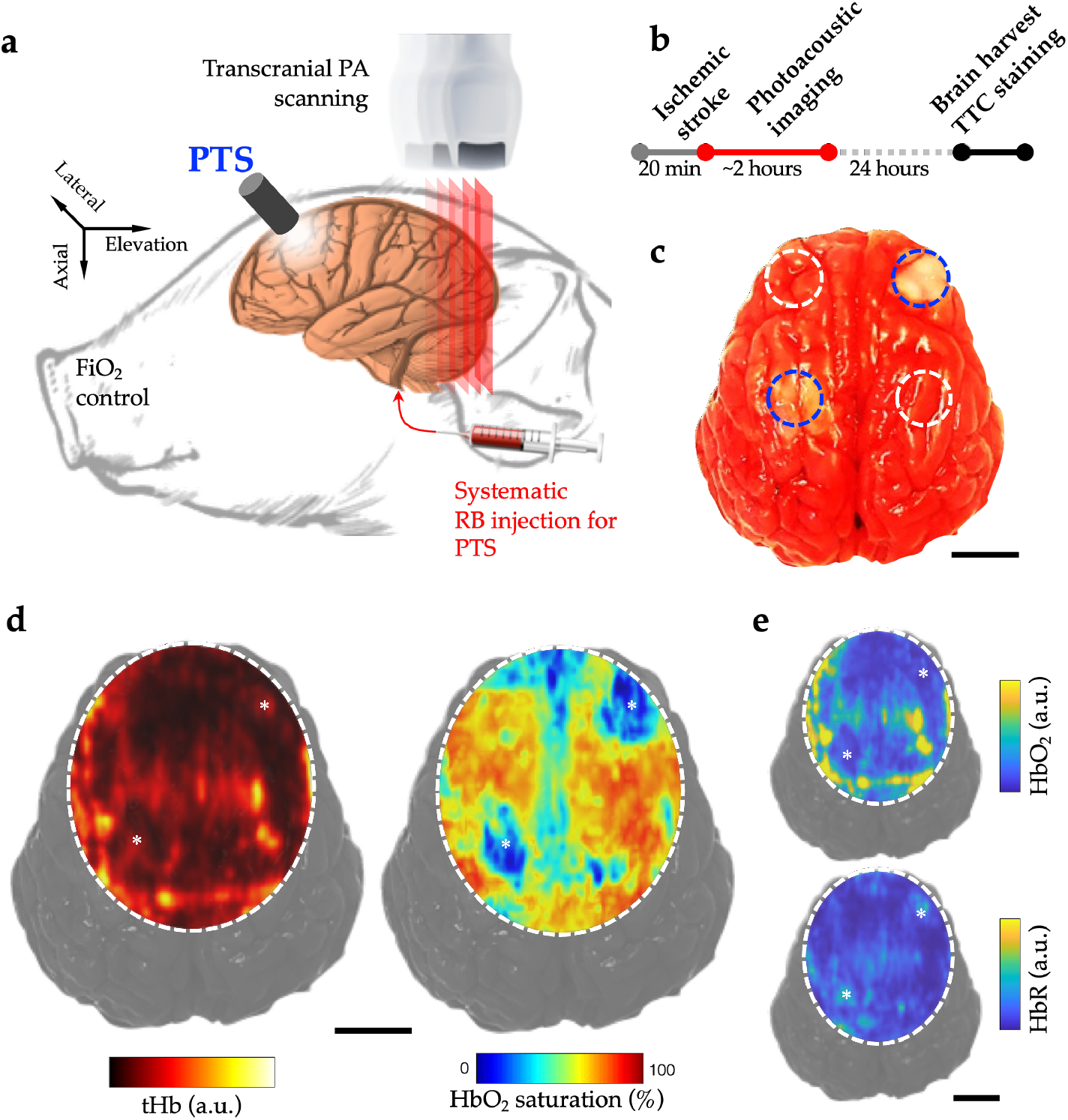
The transcranial PA imaging of photothrombotic stroke in neonatal piglets *in vivo*. (a) Experimental setup and (b) protocol for PTS induction and PA imaging. (c) Triphenyltetrazolium chloride (TTC) image of harvested whole brain, indicating cerebral infarction developed at 24-hr post-stroke time point. Blue and white dotted circles indicate the PTS lesions and control regions-of-interest. (d) Representative transverse planes of HbO_2_ saturation, total hemoglobin (tHb) overlaid on the TTC image. (e) Corresponding transverse maps of HbO_2_ and HbR quantities. Black bars indicate 1 cm. Asterisk marks indicate the cortical regions presented cerebral infarction in TTC image.

### Transcranial PA imaging of PTS lesions in neonatal piglet *in vivo*

Figures 1a and 1b show the experimental setup and protocol for the induction of regional ischemic stroke and transcranial PA imaging within 2 hours, followed by a triphenyltetrazolium chloride (TTC) staining to confirm eventual development of cortical infarction at 24 hours from the stroke induction. Figure 1c shows a representative TTC image, illustrating clear development of infarction via PTS technique. Dotted circles indicate the candidate PTS regions, and blue and white colors indicate the affected and control regions-of-interest (ROIs). Apparent regional developments of cerebral infarction were identified at the regions where the PTS protocol was applied (blue dotted circles), compared to the control ROIs (white dotted circles). Figure 1d shows the representative transcranial PA imaging in neonatal piglet *in vivo* obtained within 2 hours from the PTS onset. The transverse image of HbO_2_ saturation and tHb shows the maximal and minimal intensity projection maps over 3-mm depth range in the superficial cortical region. In particular, HbO_2_ saturation clearly localized PTS lesions corresponding to where they manifested in the TTC image (Figure 1c). Figure 1e presents HbO_2_ and HbR distributions in the corresponding transverse PA image. In visual assessment, interestingly, low HbO_2_ and higher HbR were introduced in PTS lesions when compared to control ROIs in the contralateral sides.

Figure 2 and Table 1 present and summarize the quantitative measurements of HbO_2_ saturation, tHb, HbO_2_, and HbR in the PTS model. In the piglets with a single lesion (Figure 2b), the HbO_2_ saturation was the most obvious metric giving robust statistical significance between control and affected ROIs (*p* < 0.0001, *n* = 5): 54.53 ± 6.16% vs. 21.19 ± 4.59% (mean ± SD). The metrics of tHb and HbO_2_ displayed trends for lower levels in the affected ROIs, but these did not attain statistically effective separation with this small sample size. The values of HbR were not different between the control side (1.63 ± 1.85 a.u.) and affected side (1.14 ± 0.67 a.u), although one outlier value was evident with high HbR on the control side of the brain. Rejection of the case with an outlier out of standard deviation still did not yield statistical significance: 0.94 ± 0.58 vs. 0.82 ± 0.45 a.u. In the piglets with the induction of two PTS lesions (Figure 2c), the HbO_2_ saturation was still the most effective single metric (60.01 ± 13.46% vs. 31.31 ± 14.43% in control and affected ROIs, respectively), although the decrease in HbO_2_ still did not attain statistical significance despite the higher sample size (12 lesions in 6 piglets). The HbR concentration in the double-lesioned piglets did display the expected increase in the affected ROIs when compared to those at the control ROIs: 1.27 ± 0.47 vs. 1.94 ± 0.58 in arbitrary unit (a.u.), respectively (*p* = 0.0045). Finally, we analyzed the pooled data to have a generalized perspective how accurate PA imaging can detect PTS lesions (Figure 2a). The HbO_2_ saturation again yielded strong statistical significance: 58.40 ± 11.86% vs. 28.33 ± 13.08% in control and affected ROIs, respectively. Notably, HbO_2_ quantity presented a statistically significance decrease with PTS induction (*p* = 0.0211). Overall, the HbR quantity did not present a statistically significant separation; however, with rejection of the one outlier control value, a significant increase was observed (*p* = 0.0187). The tHb concentration for the pooled data did not convey statistical significance.

**Figure 2.**
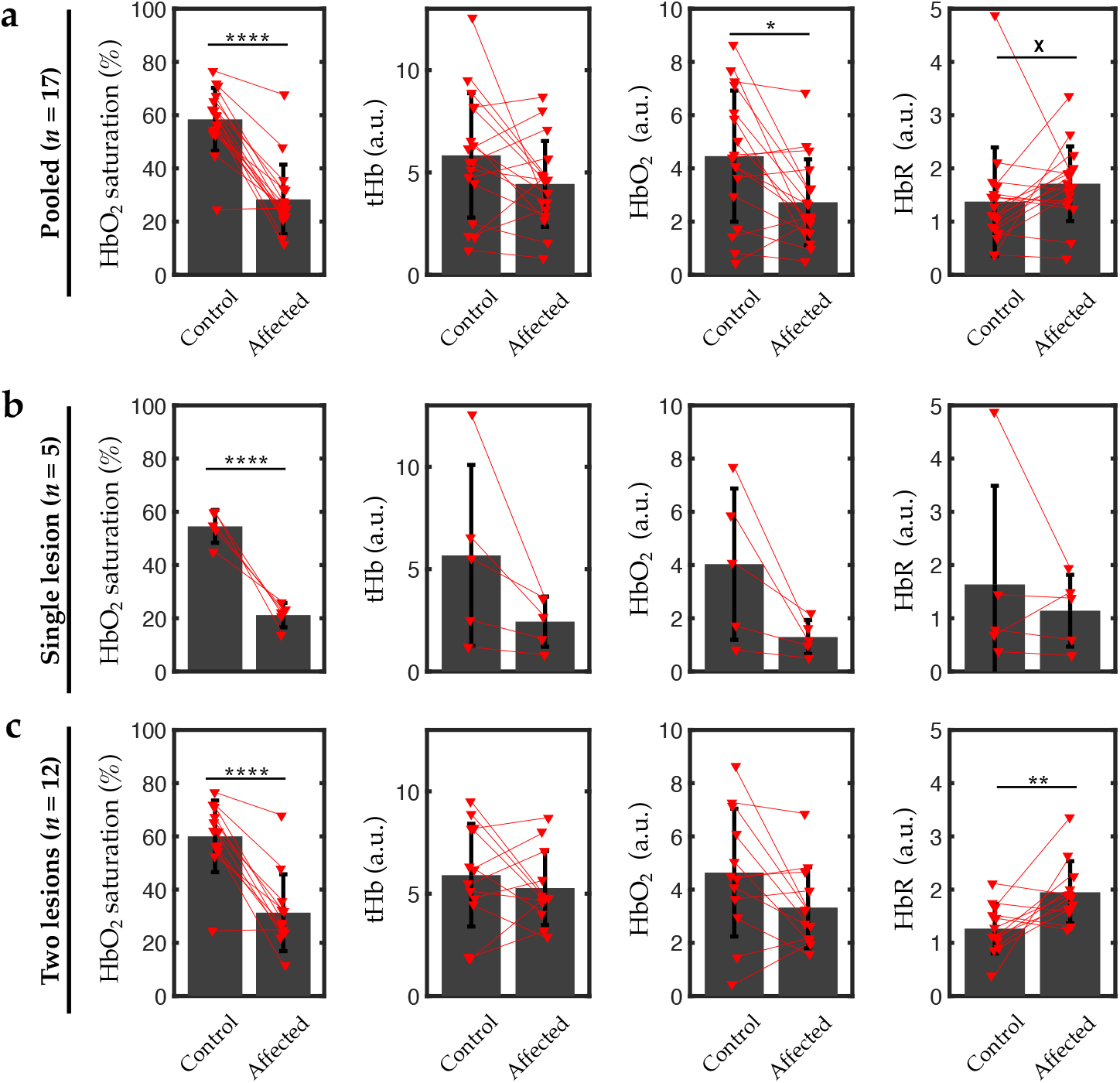
Quantitative measurements of PA imaging metrics, HbO_2_ saturation, total hemoglobin (tHb), oxyhemoglobin (HbO_2_), and deoxyhemoglobin (HbR). (a) Pooled group (*n* = 17). (b) Single-lesion group (*n* = 5). (c) Two-lesion group (*n* = 12). Red indicators show the individual data points from the regions-of-interest (ROI). Red lines indicate the control and affected ROIs from individual piglets. Asterisk marks indicate the degree of statistical significance in paired t test: * (*p* ≤ 0.05), ** (*p* ≤ 0.01), *** (*p* ≤ 0.001), and **** (*p* ≤ 0.0001). ☓ indicates a statistical significance in a specific case when removed one outlier animal data (*n* = 16, *p* ≤ 0.05).

**Table 1.** Photoacoustic (PA) imaging metrics of photothrombotic (PTS) model *in vivo*. Asterisk marks indicate the statistical significance between the control and affected regions-of-interest (ROIs). Asterisk marks indicate statistical significance: * (*p* ≤ 0.05), ** (*p* ≤ 0.01), *** (*p* ≤ 0.001), and **** (*p* ≤ 0.0001).

| Pooled data ($n = 17$ from 11 animals) | | | | |
| --- | --- | --- | --- | --- |
|  | HbO<sub>2</sub> saturation (%) | Total Hb (tHb) concentration (a.u.) | HbO<sub>2</sub> concentration (a.u.) | HbR concentration (a.u.) |
| Control | 58.40 ± 11.86 | 5.83 ± 3.04 | 4.46 ± 2.46 | 1.38 ± 1.02 |
| Affected | 28.33 ± 13.08 | 4.44 ± 2.10 | 2.72 ± 1.61 | 1.71 ± 0.70 |
| $p$ | < 0.0001 (****) | 0.0733 | 0.0066 (*) | 0.2314 ($p = 0.0118$ when $n = 16$ excluding an outlier, *) |
| AUC | 0.9204 | 0.6540 | 0.7093 | 0.7128 |
| Single PTS lesion ($n = 5$ from 5 animals) | | | | |
| Control | 54.53 ± 6.16 | 5.67 ± 4.42 | 4.03 ± 2.84 | 1.63 ± 1.85 |
| Affected | 21.19 ± 4.59 | 2.44 ± 1.22 | 1.29 ± 0.64 | 1.14 ± 0.67 |
| $p$ | 0.0019 (**) | 0.1070 | 0.0720 | 0.4817 |
| AUC | 1.0000 | 0.7200 | 0.8000 | 0.5200 |
| Two PTS lesions ($n = 12$ from 6 animals) | | | | |
| Control | 60.01 ± 13.46 | 5.90 ± 2.51 | 4.64 ± 2.40 | 1.27 ± 0.47 |
| Affected | 31.31 ± 14.43 | 5.27 ± 1.80 | 3.32 ± 1.52 | 1.94 ± 0.58 |
| $p$ | < 0.0001 (****) | 0.4103 | 0.0586 | 0.0128 (**) |
| AUC | 0.8889 | 0.6111 | 0.6875 | 0.8403 |

Figure 3 shows the critical threshold for the PA imaging metrics to identify PTS lesions from control ROIs. Note that the solid and dotted black lines indicate the sensitivity and specificity, and the blue line indicates the accuracy. First of all, HbO_2_ saturation yielded effective separation between control and affected ROIs (Figure 3a), as expected from the clear statistical separation seen in Figure 2. We here define the effective threshold range giving sensitivity and specificity higher than 80% at the same time. In the pooled data (*n* = 17), HbO_2_ saturation successfully identified PTS lesions at an effective threshold range between 32.48 % and 53.19%; the maximal accuracy was 91.18%. Also, the PTS induction with single lesion presented with a wide effective threshold range between 23.31% and 53.91% (*n* = 5). Encouragingly, the maximal accuracy at 100 % was achieved between 25.95 % and 44.89 % of threshold range. Introduction of two PTS lesions showed a somewhat narrower effective threshold range between 36.55 % and 54.40 % (*n* = 12) with the maximal accuracy of 91.67%. Figure 3b shows the corresponding analysis on tHb concentration threshold. In general, there were no cases giving an effective threshold range. Even though the maximal accuracy of 80% was achieved with the single PTS lesion cases in the range between 3.57 and 5.50 a.u., the sensitivity in that range only reached 60%. The maximal accuracy was 66.67% in the cases with two PTS lesions. Pooled data also showed low accuracy at 70.59%. The individual analysis on HbO_2_ and HbR quantities presented moderate elevation of their accuracies (Figure 3c) but still did not yield meaningful improvement compared to HbO_2_ saturation. The maximal accuracy ranged from 60% - 80%, and very narrow effective threshold ranges were identified only with HbO_2_ quantity with single-lesion case (1.62 – 1.72 a.u).

**Figure 3.**
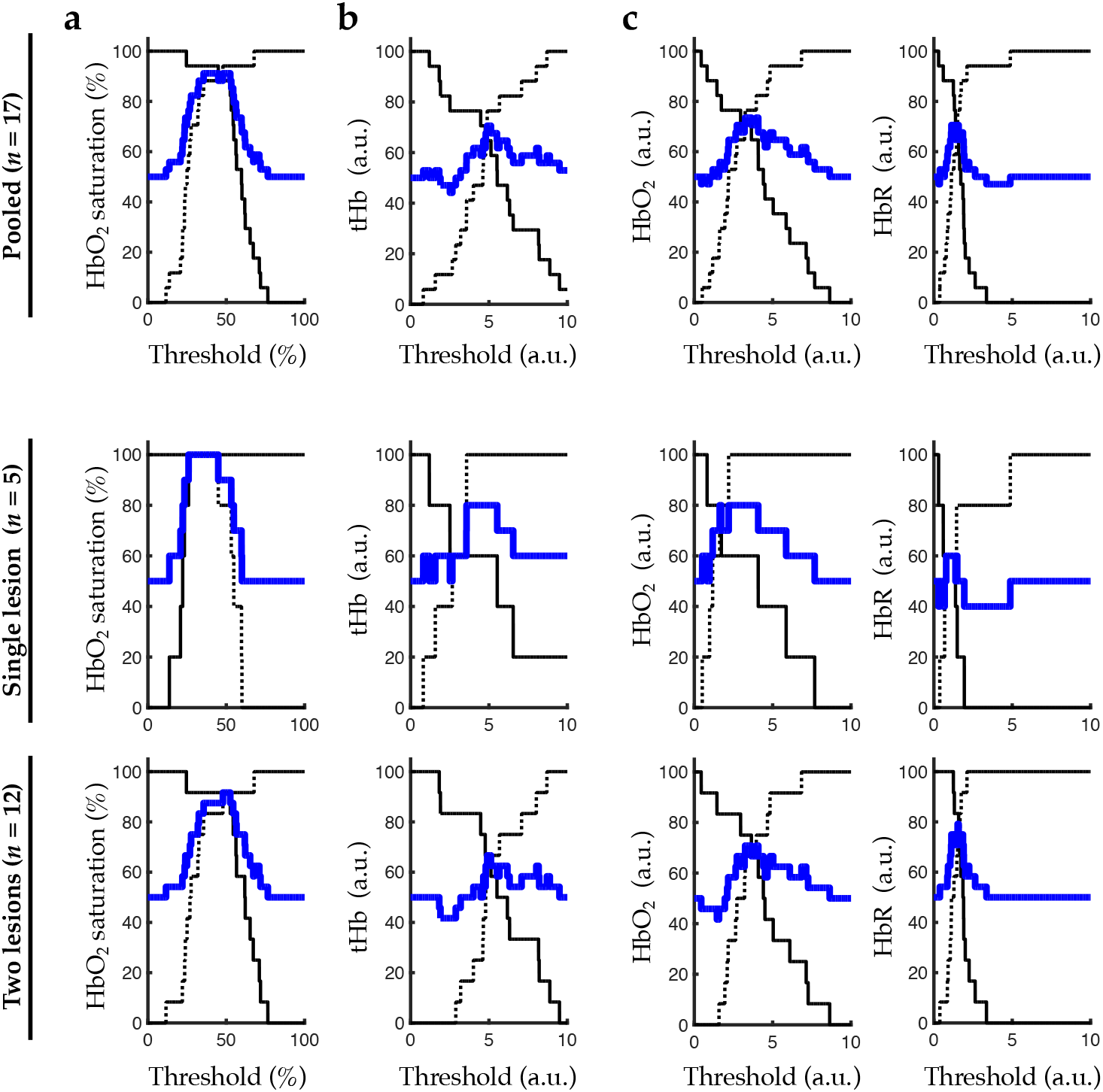
Sensitivity, specificity, and accuracy to classify the photothrombotic (PTS) lesions for the varying threshold values. (a) HbO_2_ saturation, (b) total hemoglobin (tHb), and (c) oxy- and deoxyhemoglobin (HbO_2_ and HbR) in pooled, single-lesion, and two-lesion groups. Solid and dotted black lines indicate specificity and sensitivity, respectively. Blue line indicates the accuracy.

Figure 4 shows the receiver operating characteristic (ROC) curves for a threshold-invariant evaluation of PA imaging on PTS lesion detection. Note that the dotted and solid black lines indicate the ROC curves of the cases with a single-lesion (*n* = 5) and two-lesions (*n* = 12), respectively. The blue line presents the ROC curve using the pooled data (*n* = 17). Table 1 also contains the area-under-the-curve (AUC) measurements from the ROC curves. The HbO_2_ saturation metric provided creditable classification of PTS lesions with AUC values at 0.92, 1.00, and 0.89 in pooled, single-lesion, and two-lesion cases, respectively. In hemoglobin quantity metrics, AUC value over 0.80 only appeared with HbO_2_ in the single-lesion cases and HbR in the two-lesion cases.

**Figure 4.**
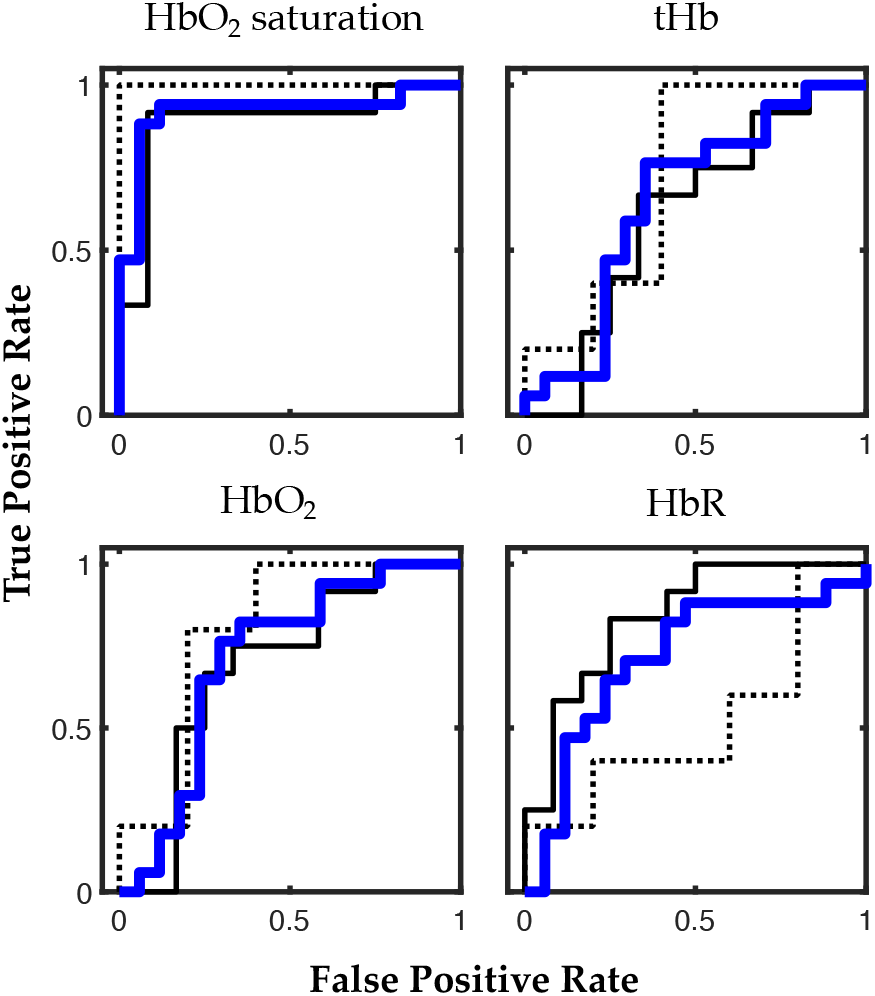
Receiver operating characteristic (ROC) curves for a threshold-invariant evaluation of PA imaging on PTS lesion detection. Blue line indicates the ROC curve with the pooled data. Dotted and solid black lines indicate the curves in the single-lesion and two-lesion groups, respectively.

### Supervised classification of PTS lesions

Table 3 summarizes the supervised machine learning-based classification accuracy on the control and affected ROIs. Firstly, a single-variable classification scenario was evaluated using individual PA imaging metrics. In general, the classification accuracy followed the statistical significance presented in Figure 3, as expected. The pooled data presented the maximal accuracy at 91.2% when only HbO_2_ saturation was included. Meanwhile, other hemoglobin quantity metrics (i.e., tHb, HbO_2_, and HbR) did not present any useful classification accuracy. The reduced data with single and two PTS lesions followed the same trends. Secondly, multivariable SVM models were analyzed with different number and combinations of variables. However, the addition of hemoglobin quantity metrics did not benefit the accuracy over the single-variable scenario solely using HbO_2_ saturation metric. The maximal accuracy in the multivariable models was 88.2% when used the pooled data and included HbO_2_ saturation as one of the variables. The cases without HbO_2_ saturation only provided up to 73.5% of accuracy. The use of other classifiers in the multivariable classification scenario (e.g., tree, discriminant, logistic regression, naïve bayes, quadratic/cubic/Gaussian SVMs, nearest neighbor, and ensemble) did not present any improvement over the single-variable classification accuracy on HbO_2_ saturation (< 91.2 %), which implies that none of them take advantage from tHb, HbO_2_ and HbR metrics.

## Discussion

In this paper, we demonstrate the feasibility of transcranial PA imaging of regional ischemic stroke in neonatal piglets *in vivo*. We found that the HbO_2_ saturation metric was sufficient for effective detection in a hand-held PA imaging system. A hand-held monitoring system has clinical translation potential for diagnosis of perinatal arterial ischemic stroke at the bedside. Moreover, the near-infrared light could penetrate the scalp and skull of the piglet and provide a PA signal with spatial resolution that was adequate for detecting an ischemic lesion on a centimeter scale. The transcranial PA imaging of HbO_2_ saturation may address limitations of other perinatal monitoring modalities: having direct cerebral hemodynamic contrast (over EFHM); continuous and prompt monitoring (over MRI); portable bedside monitoring (over MRI); and sub-millimeter-scale spatial specificity (over biophotonic modalities). However, the tissue quantity of total hemoglobin did not provide additional effectiveness in the context of the PTS model. There are many foreseeable variables in hemoglobin quantity; for example, inconsistent distance and angle of the PA probe from the fetal head would affect light fluence and acoustic attenuation. These will directly affect the estimation of hemoglobin quantity in individual patients, leading to patient-to-patient variability. This limitation could be mitigated by assuming that the ischemia is limited to a focal region of brain, thereby providing a within-patient contrast with non-ischemic tissue.

It should be noted that the PTS model used here has specific hemodynamic consequences that differ from embolic models. In particular, the measures of hemoglobin quantity suggested complex physiologic features within the affected ROIs. The affected ROIs exhibited inconsistent decreases in tHb, with as the decreases inHbO_2_ coincided with increases in HbR. Thrombosis evoked by endothelial damage from the light/dye interaction likely starts in venules and capillaries, where shear stress is lower than in arterioles. Furthermore, because venules and capillaries have no smooth muscle layer, damaging their endothelium will more readily expose underlying collagen in the extracellular matrix, which activates platelets and causes leukocytes to stick to the endothelium. The resulting thrombosis will cause entrapment of red blood cells within the microcirculation. This entrapment may account for the inconsistent decrease in tHb reported by PA imaging in this model. In contrast, embolic occlusion of a major cerebral artery reduces inflow into the microcirculation but does not block outflow. Indeed, PET images of brain in adult ischemic stroke show a decrease in cerebral blood volume that correlates with the decrease in cerebral blood flow [36,37]. Thus, the tHb response may differ with a perinatal stroke in humans, in which a clot typically occludes large cerebral arteries rather than vessels in the microcirculation. Unfortunately, the rete mirabilis between the pharyngeal artery and the intracranial internal carotid artery in the pig does not permit induction of a focal stroke in only a portion of the cerebral hemisphere by insertion of a blunted tip filament or blood clot or by embolization with microspheres. Therefore, further investigation of PA imaging capabilities in a model of embolic stroke in the piglet would likely require a full craniotomy to directly occlude a major cerebral artery.

Several engineering strategies could be employed to improve and supplement the proposed framework. The spatial resolution of the transcranial PA imaging is subject to the degree of phase aberration when imaging through skull layers, having heterogenous thickness with sound propagation speeds significantly different from that of soft biological tissue (4,080 m/s vs. 1,540 m/s). Even though the human newborn skull is thinner than adults and more transparent at the open fontanelles, the use of effective phase aberration correction will further improve the spatial resolution [38]. In addition, spectral attenuation in scalp and skull layers also needs to be corrected for accurate spectral unmixing [39]. ‘Spectral coloring’ artifacts are generated by light absorbance at fetal scalp and superficial vessels containing abundant melanin and blood, resulting in lower spectral unmixing accuracy due to the discrepancy between a measured PA spectrum and reference absorption spectrum of hemoglobin. Even though we compensated the artifacts with *ex vivo* calibration, more advanced fluence estimation and correction at individual imaging wavelengths will further improve the imaging quality for accurate stroke identification.

Our long-term goal is to transform our transcranial PA imaging system into a safe, compact, and cost-effective form factor to facilitate its clinical translation. The investigation should be based on promising progress in compact light source (pulse laser diode [40], light-emitting diode [25], etc.) and ultra-portable ultrasound imaging systems [41–43]. For example, monitoring of a vulnerable neonate may also necessitate a compact yet effective monitoring device integrated in the neonatal intensive care unit (NICU). The careful tradeoff between image quality and compactness for fluent workflow would be a key for a successful clinical translation.

In adult stroke, early imaging with CT or MRI provides a differential diagnosis of ischemic versus hemorrhagic stroke and selecting patients for treatment with tissue plasminogen activator and endovascular thrombectomy [44–46]. Early imaging results also allows for selection of patients in clinical trials that are most likely to benefit from therapeutic interventions. In contrast, clinical trials of newborn stroke are hampered by constraints of imaging unstable newborns in the first few hours after birth and the limited ability to differentiate intrapartum global hypoxia-ischemia from a focal stroke based on the early neurological exam. Newborns with stroke can present with seizures by 12 hours after birth [11], but neuroprotectants are unlikely to have a major benefit when initiated after seizure onset. Therefore, the ability to diagnose a focal stroke within a few hours after birth could have a major impact by enabling early enrollment and stratification in clinical trials for early treatments. Hence, noninvasive PA imaging, by differentiating focal stroke from global hypoxia-ischemia, has great potential for advancing clinical trials of neonatal stroke, along with promising technological progresses [47,48].

In conclusion, a transcranial PA imaging is capable of identifying a focal thrombotic stroke in a neonatal piglet model with intact scalp and skull at an early stage before the infarction is formed. Furthermore, the PA-derived HbO_2_ saturation is sufficient for providing robust sensitivity and specificity for detecting 1 cm focal cortical strokes. This technology has great potential for rapid and early diagnosis of perinatal ischemic stroke in clinics.

## Materials and Methods

### Photothrombotic model in neonatal piglets *in vivo*

All procedures were approved by the Johns Hopkins University Animal Care and Use Committee. Aseptic surgery was performed on male piglets anesthetized with isoflurane (approximately 1.5%) during the whole procedure. The position of piglet head was stabilized by using a custom stereotaxic equipment. The lungs were ventilated via intubation (endotracheal tube Tube, 2.5 mm, cuffed, Medsource Labs, MN, USA) with approximately 25% O_2_ and 75% air. A femoral artery and a jugular vein were catheterized, and the mean arterial blood pressure was continuously monitored. Rectal temperature was maintained near 38.0~38.5 °C with a warm blanket. Arterial pH, PCO_2_, PO_2_, Hb, HbO_2_ saturation, and glucose were measured with a blood gas analyzer (ABL800 FLEX blood gas analyzer, Radiometer America Inc. Brea, CA, USA). A fluid of 5% dextrose and 0.45% sodium chloride was infused through jugular vein at the rate of 10 ml/h. Eleven male piglets, weighing 1.57 ± 0.18 kg were included in this study, with 5 in the single-lesion and 6 in the two-lesion PTS groups. Their mean arterial blood pressure at baseline was 61.3 ± 12.8 mmHg, and mean temperature was at 37.5 ± 0.7 oC. Blood sample data are summarized in Table 2. Note that we did not make any attempt to study equal numbers of male and female piglets because we are not aware of any plausible biological reason for sex differences in neurovascular physiology for direct photothrombosis.

**Table 2.** The arterial blood sample analysis of the piglets during stroke induction experiment (Data format: Mean ± SD). Abbreviations: pCO_2_, Arterial carbon dioxide partial pressure; pO_2_, Arterial oxygen partial pressure;

| pH | pCO<sub>2</sub> (mmHg) | pO<sub>2</sub> (mmHg) | cBase (mmol/L) | HbO<sub>2</sub> saturation (%) | ctHb (g/dL) |
| --- | --- | --- | --- | --- | --- |
| 7.37 ± 0.09 | 37.1 ± 5.5 | 143.5 ± 23.6 | -3.5 ± 3.3 | 99.7% ± 0.5% | 11.0 ± 2.9 |
| K<sup>+</sup> (mmol/L) | Na<sup>+</sup> (mmol/L) | Ca<sup>2+</sup> (mmol/L) | Cl<sup>-</sup> (mmol/L) | Glu (mg/dL) | Lac (mmol/L) |
| 3.7 ± 0.6 | 141.9 ± 3.4 | 1.3 ± 0.2 | 108.7 ± 5.1 | 141.9 ± 49.6 | 4.5 ± 8.6 |

**Table 3.**
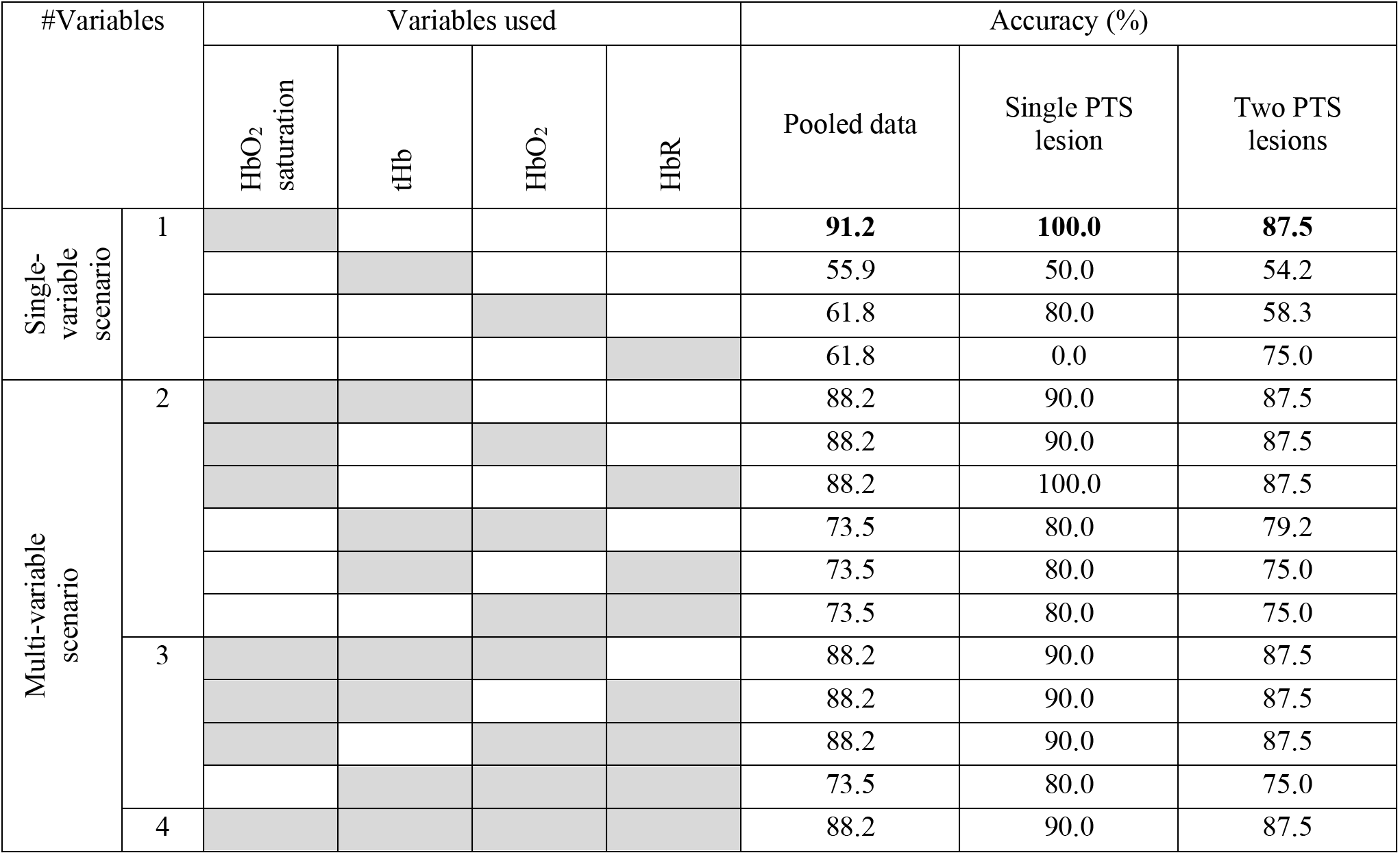
Supervised machine learning-based differential classification accuracy using a linear support vector machine (SVM) classifier. 34 data points (17 ROIs for both controlled and PTS lesions) were used with 34-fold cross-validation. Gray shadings indicate the variables used.

A solution of 20 mg/kg Rose Bengal (4,5,6,7-tetrachloro-2’,4’,5’,7’-tetraiodofluorescein, RB) s in sterile saline was injected via jugular vein. The dye has an absorbance peak at approximately 560 nm [49], and the skull was thinned with a drill to allow sufficient light to penetrate the cortex. The focal skull areas overlying the frontal (5 mm anterior and 8 mm lateral from bregma) and posterior brain tissue regions (5 mm posterior and 8 mm lateral from bregma) was thinned with 5-mm width. The skull thinning was performed bilaterally, but only half of the focal areas were targeted for photothrombosis. In those targeted for photothrombosis, the thinned skull was illuminated with a 5-mm aperture cold-white light at the intensity of 60 – 75 mW/cm^2^ for 20 minutes. The light interaction with the dye generates free radicals that damage the endothelium sufficiently to activate platelets and produce a thrombus. During the light illumination, 1 ml sterile saline solution was perfused on the thinned skull every 5 minutes to prevent overheating and damage on target surface. No signs of thermal damage due to the light illumination were found with light illumination in the absence of Rose Bengal dye with regular saline flushing.

### Photoacoustic imaging system and protocol

Figure 1a shows the experimental system configuration. For PA recording, 5.2-MHz clinical ultrasound linear array transducer (L7-4, ATL Ultrasound Inc.) was connected to a research package (Vantage 256, Verasonics Inc., USA). Detailed imaging parameters are as follows: elevation focal depth, 25 mm; element pitch, 0.30 mm; element height, 7.5 mm for transducer, 18 and 24 dB low-noise amplification (LNA) settings during ultrasound (US) and PA imaging. Those parameters and system configuration are available in modern clinical US imaging platforms, maximizing the translationability of the research outcome into clinics. To induce the PA signals, pulsed laser light was generated by a second-harmonic (532 nm) Nd: YAG laser pumping an optical parametric oscillator (OPO) system (Phocus Inline, Opotek Inc., USA). The tunable range is 690-900 nm and the maximum pulse repetition frequency is 20 Hz. Light was delivered into the probe through bifurcated fiber optic bundles, each with 40-mm long and 0.88-mm wide aperture. The PA probe is located on the center of piglet head along with the coronal plane to simultaneously monitor the PA signal changes originated from superior sagittal sinus and cerebral brain tissue. The distance between fiber bundle output and piglet head was maintained at 20 mm, leading to the 5 mJ/cm^2^ of energy density on the skin surface. This is far below the maximum permissible exposure (MPE) of skin to laser radiation guided by the ANSI Z136.1 standard, Safe Use of Lasers (i.e., 20 mJ/cm^2^). [54] In addition, 64 subsequent frames were averaged to overcome the limited signal-to-noise ratio (SNR) at cerebral brain tissue regions with naturally less blood content than the sagittal sinus and to alleviate the energy fluctuation in the Nd:YAG OPO laser source.

Figure 1b shows *in vivo* experimental protocol. At 20-min after inducing the last PTS, translational scanning was initiated from frontal to posterior regions for volumetric PA imaging within 20-30 min. The translational scanning was performed from front to the back of the brain in 1 mm intervals with a motorized stage (MTS50/M-Z8 and TDC001, Thorlabs Inc., USA). The HbO_2_ saturation and tHb were decomposed from spectral PA data (700–900 nm in 10 nm interval), and transverse planes were projected in axial direction at every 1 mm depth interval. From the data, spectral unmixing was conducted by a constrained linear least-squares problem solver in MATLAB software (Mathworks Inc., USA). Optical absorbance of HbO_2_ and deoxy-Hb was obtained from spectrophotometric measurement (brought from [50]). The mean spectral attenuation was measured *ex vivo* and compensated before the spectral unmixing process [24].

### Triphenyltetrazolium chloride (TTC) staining and imaging

After the PA scanning, the isoflurane was stopped, and the piglet was carefully removed from the stereotaxic apparatus. The piglet was kept on a pre-warmed heating pad until it was fully awake The pig was extubated when it regained a cough reflex.. The piglet was returned to the cage and fed formula milk. On the next day, the piglet was euthanized with an overdose of pentobarbial, and the brain was removed for staining with the vital dye TTC (2% solution) on fresh tissue. TTC stains intact mitochondrial enzymes red whereas the infarcted region in pale (Figure 1c). The surface contour of cerebral infarction carefully was drawn by one of the authors (X. Liu). We correlated the PA images with 24-hr post-stroke focal infarction developments on cortex detected through TTC image.

### Statistical analysis

PA metrics were compared between the affected ROIs and the contralateral control ROIs by paired t-test. ROC curves and corresponding AUC were reconstructed and calculated in MATLAB software (R2020a, Mathworks, Inc., USA) using *perfcurve* function.

### Supervised machine learning

The pooled measurements from control and affected ROIs (*n* = 34) were fed into a supervised machine learning classification learner. Various combinations of PA imaging metrics (i.e., HbO_2_ saturation, tHb, HbO_2_, and HbR) were trained in a linear SVM model using MATLAB software (R2020a, Mathworks, Inc., USA), and tested with 34-fold cross validation scheme (i.e., 33 training data and 1 test data; repeat 34 times).

## Acknowledgements

The financial support was provided by NIH, NHLBI (R01HL139543); Maryland Innovation Initiative, TEDCO; Louis B. Thalheimer Fund for Translational Research.

